# U2-2 snRNA Mutations Alter the Transcriptome

**DOI:** 10.1101/2025.03.12.642709

**Authors:** Zhongxuan Chi, Varun Gupta, Charles Query

## Abstract

Intron removal from pre-mRNA is catalyzed by the spliceosome, which comprises 5 snRNPs containing small nuclear RNAs (snRNAs). U2 snRNA makes critical RNA-RNA and RNA-protein contacts throughout the splicing cycle. Mutations in U2 snRNA, particularly at position C28, have been linked to cancers. To study gene expression changes mediated by mutated U2 snRNAs, U2-2 C28 mutants, U2-2 knockout (KO), and U2-2 overexpression (OE) cell lines were constructed followed by RNA sequencing. We observed significant changes in splicing and over 4,000 differentially expressed genes enriched in pathways like RNA processing and non-coding RNAs upon knocking out U2-2 snRNA. Splicing patterns were more influenced by U2-2 dosage than mutations alone. Therefore, the mutant exhibits a compound phenotype, resulting from reduced U2-2 levels (and thus mostly phenocopying the KO) and additional mutant-specific splicing changes.

**HIGHLIGHTS:**

- U2-2 snRNA BSL mutants alter splicing and the transcriptome
- U2-2 KO phenocopies most altered splice events in the mutants
- Both U2-2 levels and mutations alter splicing
- Many altered splice events lead to NMD

## INTRODUCTION

Alternative pre-mRNA splicing is a fundamental part of gene expression that drives transcript diversity (Wang et al., 2008) and is crucial for numerous biological processes, including development and cell-type specification (Chen & Manley, 2009; Ellis et al., 2012; Graveley, 2001; Kalsotra & Cooper, 2011; Pan et al., 2008; Yang et al., 2016). Through the precise removal of introns, splicing converts long precursor pre-mRNAs into mature mRNA (Han et al., 2011). This complex process is catalyzed by the spliceosome, a dynamic ribonucleoprotein complex composed of five small nuclear ribonucleoproteins (snRNPs), which include the small nuclear RNAs (snRNAs) U1, U2, U4, U5, and U6, along with hundreds of proteins (Lamond, 1993; reviewed in Wahl et al., 2009; Will & Lührmann, 2011). U2 snRNA plays an essential role in splicing, as it forms critical RNA-RNA and RNA-protein contacts during the splicing cycle. Specifically, the branchpoint-interacting stem loop (BSL) region of U2 pairs with the branch site region during initial steps of splicing (Parker et al., 1987; Perriman & Ares, 2010). The BSL of U2 snRNA is evolutionarily conserved across species, indicating that the BSL is an ancient structure of intron branchpoint recognition (Perriman & Ares, 2010). Additionally, mutations in the BSL alter splicing in *S. cerevisiae* (Perriman & Ares, 2010). The ATPase Prp5, required for stable U2 snRNP binding to the intron, is postulated to destabilize the BSL by direct interaction with the BSL RNA and/or by removing Cus2/Tat-SF1, allowing for branch region base pairing and providing a platform for incoming U6 snRNA during spliceosome assembly (reviewed in van der Feltz and Hoskins, 2019). Both weakened BSL and hyperactive *prp5* ATPase mutants lead to increased branch recognition fidelity, whereas strengthening the BSL or slowed *prp5* ATPase mutants lead to decreased fidelity (Xu & Query, 2007). Mutations in the BSL can rescue *prp5* mutants (Perriman and Ares, 2010), and evidence suggests that the identity of some BSL nucleotides can alter Prp5p ATPase activity (Wu et al., 2016). Likewise, some mutations in the BSL disrupt base pairs formed with U6 snRNA (helix Ia and helix III) in the full spliceosome; these changes also impair splicing and can be rescued by compensatory changes in U6 (Madhani and Guthrie, 1992; Ryan & Abelson, 2002; Smith et al., 2009). Thus, the identify of U2 BSL nucleotides can play crucial roles, perhaps at multiple stages in spliceosome assembly and later spliceosome function to affect the outcome of splicing.

In humans, there are over 100 U1, more than 72 U2, and over 1300 U6 gene loci. However, with few exceptions [e.g. (Cheng et al., 1997; Jia et al., 2012; Lund et al., 1984, 1985) and the 6 abundant U5s (Sontheimer & Steitz, 1992)], only canonical snRNAs have been studied. Canonical U2 snRNAs are highly conserved and crucial for splicing across species. The two most abundant U2 snRNAs, RNU2-1 (U2-1) and RNU2-2 (U2-2), are expressed in tissue-specific ratios, with U2-2 being particularly abundant in the cerebral cortex (Kosmyna et al., 2020). U2-1 snRNA is encoded by a multicopy array, the copy number variation (CNV) of U2-1 was detected by combed DNA/FISH analyses, where each of the two alleles ranged from 6-82 copies (Lindgren et al., 1985; Tessereau et al., 2013). Whereas the repetitive structure of U2-1 complicates genetic manipulation in studies, U2-2 is encoded by a single locus on chromosome 11 that is easily altered genetically (Kosmyna et al., 2020), and we show here that HEK293T U2-2 KO cells generated by CRISPR/Cas9 are viable.

The role of altered splicing in cancer has recently received much attention (Grosso et al., 2008; Sveen et al., 2016). Many studies have reported that mutations in splicing components correlate with high frequency of cancers (Agrawal et al., 2017; Bousquets-Muñoz et al., 2022; Cooper et al., 2009; Darman et al., 2015; Jia et al., 2012; Licatalosi & Darnell, 2006; Shuai et al., 2019; Suzuki et al., 2019). The Cancer Genome Atlas (TCGA) has identified 119 splicing factor genes with significant non-silent mutation patterns across 33 tumor types. Other examples include mutations in multiple splicing components, including six ubiquitous core snRNP proteins, that cause one of the most common hereditary diseases, retinitis pigmentosa (RP); there are more than 70 genes identified with altered splice events, and most of these genes are specifically expressed in the retina (reviewed by Růžičková & Staněk, 2017). Additionally, SF3b1 is one of the most mutated spliceosomal components in many diseases (Agrawal et al., 2017; Biankin et al., 2012; Darman et al., 2015; Malcovati et al., 2015). SF3B1 disease-associated mutations induce cryptic 3’ss selection by using a different branch site in splicing. Furthermore, novel SF3B1 in-frame deletions have been associated with chronic lymphocytic leukemia (CLL) (Agrawal et al., 2017). Although there are many mutations of protein-coding factors in splicing being extensively studied, little is known of mutations in snRNA components of the spliceosome. One known example of an snRNA linked to disease is U4atac snRNA, mutations of which give rise to developmental disorders such as microcephalic osteodysplastic primordial dwarfism type 1 (MOPD1) or Taybi-Linder syndrome (TALS)(Benoit-Pilven et al., 2020). A second is the U1 snRNA, recurrently mutated in multiple cancers (Shuai et al., 2019); a recurring A>C somatic mutation at the third base of U1 snRNA disrupts global gene splicing and expression in patients with chronic lymphocytic leukemia (CLL) and hepatocellular carcinoma (HCC). Another example of U1 snRNA being linked to diseases is the association between the reduction of U1 snRNP and Alzheimer’s disease (AD)(Bai et al., 2013; Campagne, 2024; Z. Cheng et al., 2021; Diner et al., 2014). Specifically, U1-70K knockdown or U1 snRNP inhibition increases amyloid precursor protein levels; this leads to abnormal RNA splicing and AD pathogenesis (Bai et al., 2013).

For U2 snRNA mutations, a mouse U2 snRNA mutation that causes neurological defects was found using a forward genetic screen (Jia et al., 2012). Challenges in identifying snRNA mutations is due to the repetitive nature of snRNA genes (Denison et al., 1981; Khurana et al., 2016; Manser & Gesteland, 1982). Large tandem repeat sequences give rise to mapping limitations and absence in reference genomes. We recently developed a new pipeline to uniquely map whole genome sequence (WGS) reads to U2 gene loci and identified across the TCGA a cluster of cancer-associated mutations at nts 25-29 in the U2 BSL (Gupta and Query, unpublished data). In addition, a recent computational pipeline, *Armadillo,* enabled the identification of somatic mutations in repetitive regions (Bousquets-Muñoz et al., 2022) and identified mutations in 73 tumors across multiple cancer types at U2 snRNA position 28, including hematological malignancies, prostate cancer, and pancreatic cancer from the ICGC PanCancer Analysis of Whole Genomes (PCAWG) and mantle cell lymphoma (MCL) projects. Collectively, this suggests that position 28 of *U2* is a candidate biomarker present in these tumors. However, the impact of U2-2 snRNA on gene expression and its specific effects on splicing are not well understood. This gap in knowledge leaves many questions unanswered regarding how U2-2 snRNA influences splicing and whether it participates in regulating alternative splicing events.

To address this gap, we generated U2-2 snRNA KO and OE cell lines and performed RNA sequencing to assess the impact of U2-2 snRNA on transcriptome-wide splicing and gene expression. Here, we found that U2 snRNA KO disrupts splicing patterns, with mutant lines showing variations in splicing severity. Over 4,000 differentially expressed genes were identified, highlighting pathways involved in ribonucleoprotein (RNP) complex biogenesis, ribosome biogenesis and rRNA processing. In addition, OE of U2-2 WT and C28T mutants further demonstrated distinct gene expression changes. Splicing patterns also changed based on the levels of U2-2 snRNA, with dosage having a stronger impact than mutations alone. Overall, these findings reveal that U2-2 snRNA plays a critical role in regulating splicing, as well as its potential as a regulatory element in cancer-associated splicing alterations.

## RESULTS

### Generation of U2-2^KO/KO^ and U2-2 mutant HEK293T cell lines

There are 6 nucleotide differences in mature U2-1 and U2-2 snRNAs (Supplemental Figure S1A), and they also differ in their promoter and 3’ splice site (3’ss) regions. The mutation C28 is within the BSL of U2 snRNA (Supplemental Figure S1B), which is important for RNA-RNA interactions with both U6 snRNA and pre-mRNA during early spliceosome assembly (Supplemental Figure S1C). Despite the structural importance of U2-2 C28, its recurrent mutations observed in many diseases are unexplained. We utilized CRISPR/Cas9 to delete the U2-2 snRNA coding sequence from the genome (Figure 1A). HEK293T cells were transfected with two plasmids, each containing Cas9, GFP, and a single guide RNA (gRNA) targeting either upstream or downstream of the U2-2 gene (“*RNU2-2P*”). Homologous recombination using template plasmids allowed for the insertion of C28T and C28A mutations, resulting in knock-in mutant lines. Transfected cells were then single cell sorted based on GFP positivity, and viable clones were expanded and genotyped. Three clonal U2-2^KO/KO^, two U2-2^WT/C28T^, two clonal U2-2^C28T/KO^ and two U2-2^C28A/KO^ biological replicates were generated. Genotyping confirmed 220 bp PCR products for the U2-2^KO/KO^ clones, indicating successful knockout (Figure 1B), with additional genotyping results for the remaining lines shown in Supplemental Figure S2. Amplicon sequencing was used to measure relative RNA levels of U2-1 and U2-2 based on their sequence differences (Supplemental Figure S1A and Figure 1C). Moreover, U2-2 levels accounted for roughly 17% of the total U2 snRNA in parental HEK293T cells, whereas U2-2 RNA was absent in the U2-2^KO/KO^ lines, as expected (Figure 1C). U2-2^KO/KO^ and other mutant lines (Supplemental Figure S2) were used as experimental groups compared with the HEK293T parental WT groups. Overall, these cell lines are utilized to investigate the molecular changes that occur due to U2-2 snRNA dysregulation.

**Figure 1.**
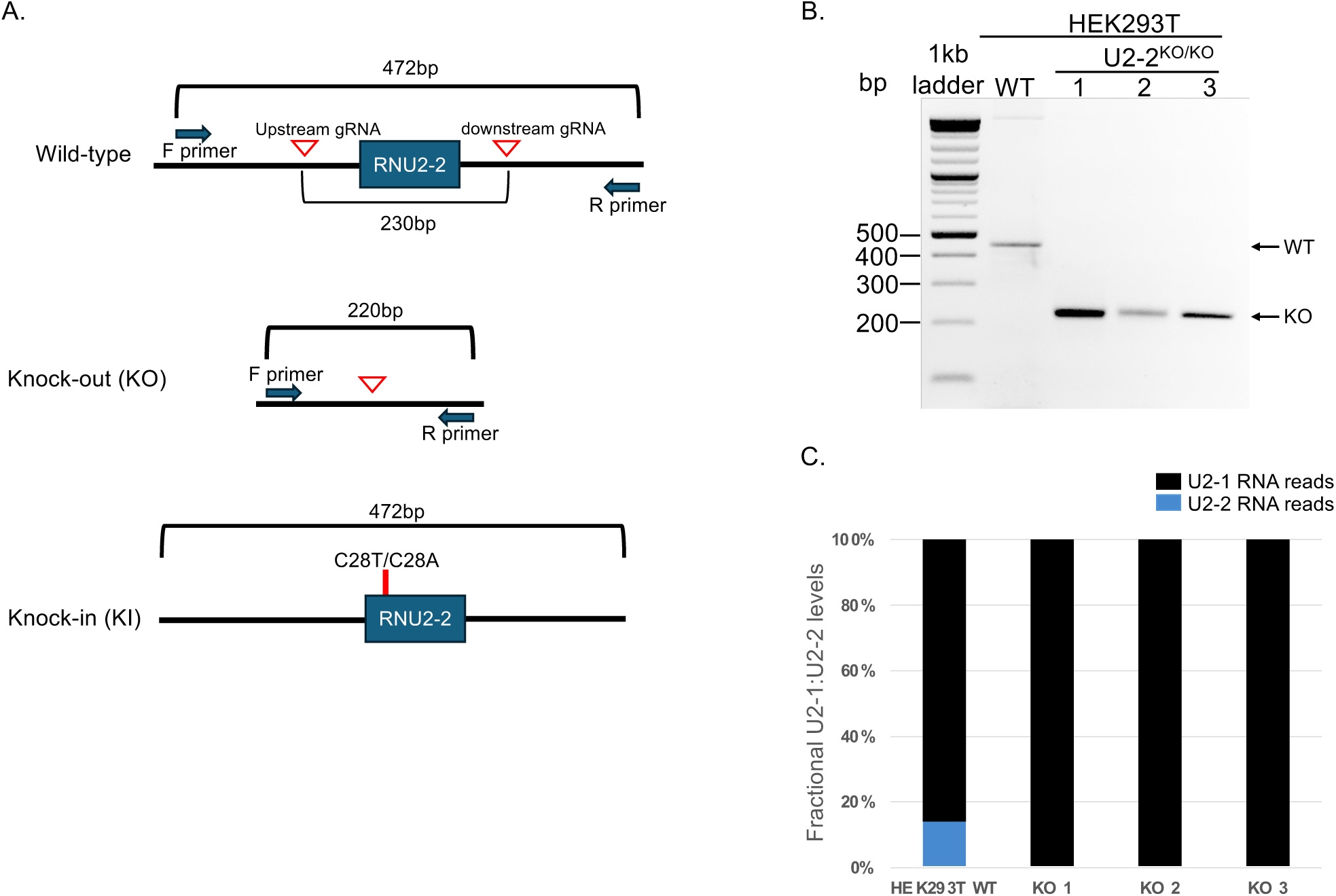
Knockout of U2-2 and knock-in of U2-2 C28T/C28A mutants in HEK293T cells. (A) Schematic representation of CRISPR/Cas9-mediated KO of the U2-2 gene and knock-in (KI) of U2-2 C28T or C28A mutants. gRNA cut sites (upstream and downstream red triangles) and genotyping primers (F and R primers, blue arrows) are indicated. (B) Agarose gel electrophoresis showing PCR products of U2-2 genomic DNA fragments from wild-type (WT) and U2-2 KO cells. The KO band (∼220 bp) is shown for three replicates, and the WT band (∼450 bp) is shown for reference. (C) Quantification of U2-2 and U2-1 snRNA levels in WT and U2-2 KO cells by RT-PCR and amplicon sequencing. U2-2 snRNA is depleted in U2-2 KO cells while U2-1 snRNA remains unchanged. Results are normalized to the total number of U2 reads. Only reads with Phred quality score q: 37 were considered. Typical read depth: ∼30,000-500,000.

### U2-2 snRNA dysregulation alters splicing and gene expression

To investigate whether U2-2 snRNA dysregulation results in transcriptomic changes, polyA+ RNA was sequenced. Five types of splice events were analyzed: skipped exons (SE), retained introns (RI), mutually exclusive exon (MXE) and alternative 5’ and 3’ splice sites (A5SS and A3SS) (Supplemental Figure S3A). Knocking out U2-2 significantly altered splicing patterns in HEK293T cells, with 2,566 SE and 611 RI significant events (FDR p-value < 0.05) between WT vs U2-2^KO/KO^ (Figure 2A and 2B). Overall splicing patterns comparing WT to U2-2^KO/KO^ and U2-2^C28T/KO^ were similar, except that U2-2^C28T/KO^ exhibited fewer MXE events (290) (Supplemental Figure S3B). A3SS and A5SS events of different comparisons are shown in Supplemental Figure S3C and S3D, respectively. Further, the U2-2^C28A/KO^ exhibited similar splicing changes as did U2-2^C28T/KO^. In contrast, there was higher inclusion of SE events in the WT compared to the U2-2^WT/C28T^, and more RI events (1,199) in U2-2^WT/C28T^ compared to WT (Figure 2A and 2B). To validate these findings, we examined the top SE events, the most prevalent splicing alteration across comparisons, using RT-PCR. We confirmed that mRNAs of *LUC7L*, *SNRNP70*, and *ZNF207* showed increased exon inclusion in the U2-2 KO and mutants compared to the WT, consistent with the RNA-sequencing results (Figure 2C-E). More importantly, these results suggest that U2-2 snRNA dosage has a greater impact on SE events than the mutant alone, as U2-2^C28T/KO^ and U2-2^KO/KO^ show more extensive splicing alterations than does U2-2^WT/C28T^ compared to the WT; furthermore, SE events in U2-2^C28T/KO^ and U2-2^KO/KO^, compared to WT (1,120), overlap more than those in U2-2^WT/C28T^ and U2-2^KO/KO^ (601), compared to WT (Figure 2E).

**Figure 2.**
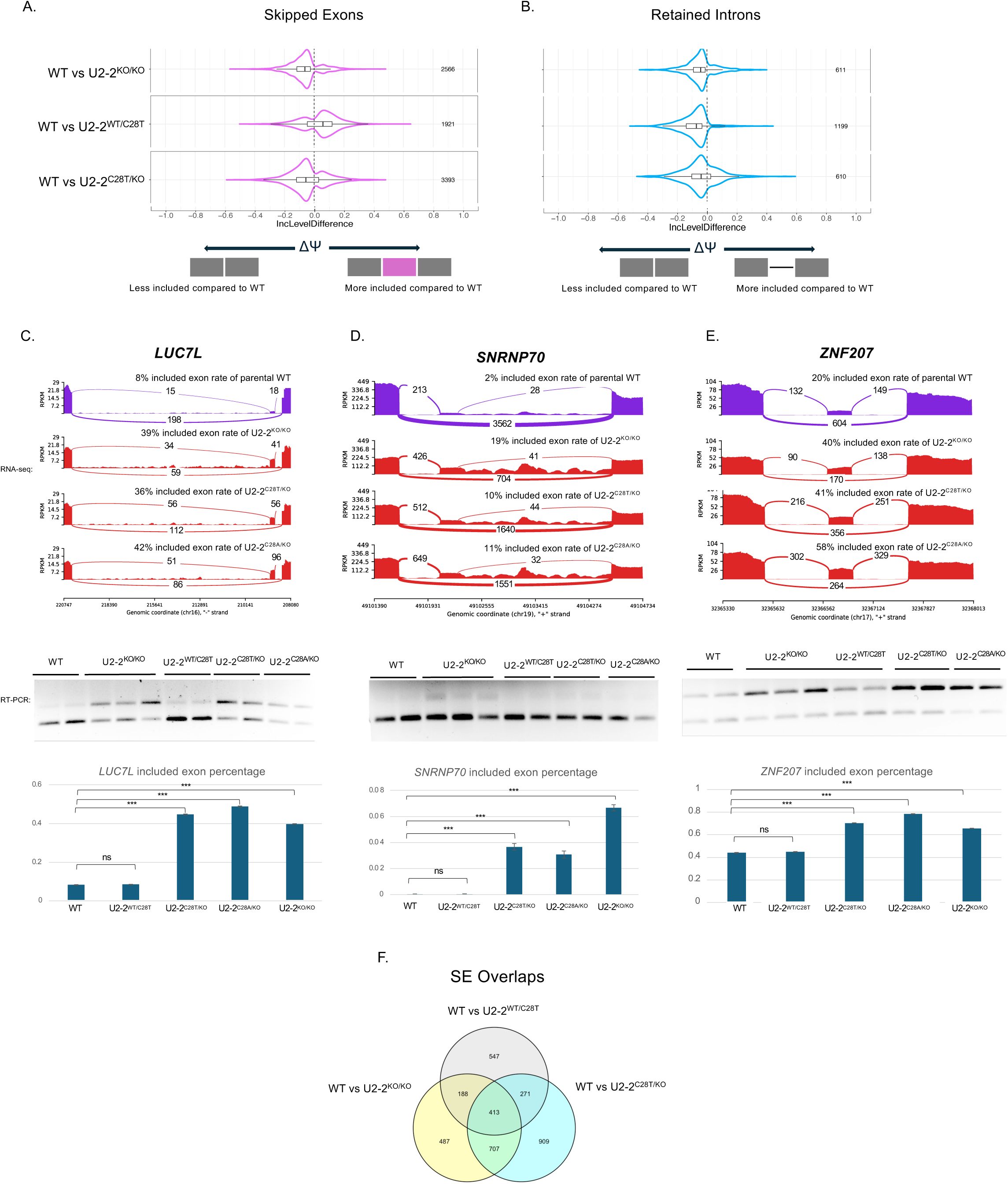
Both U2-2 KO and C28T mutants alter the splicing compared to the WT of HEK293T. (A) Comparison of ΔΨ of SE events between WT vs three U2-2^KO/KO^, two U2-2^WT/C28T^, and two U2-2^C28T/KO^ cell lines at p ≤ 0.05 cut-off. The total number of significant altered events are shown on the right. Grey boxes indicate exons and lines indicate introns. Purple boxes indicate mis-spliced exons corresponding to the SE event. (B) Comparison of ΔΨ of RI events between WT vs three U2-2^KO/KO^, two U2-2^WT/C28T^, and two U2-2^C28T/KO^ cell lines at p ≤ 0.05 cut-off. (C-D) Sashimi plots (upper) of altered SE events of *LUC7L*, *SNRNP70* and *ZNF207*, respectively, in different conditions through RNA-sequencing results. The Reads Per Kilobase per Million mapped reads (RPKMs) for each splice site are shown on the y-axis. Lower: RT-PCR was used to verify the exon inclusion percentage in different groups. cDNA from cells expressing the indicated ORFs was subjected to PCR amplification using primers flanking the alternative exons (Supplemental Table S1). PCR products were separated in a 2% agarose gel. The ratios of the intensity of PCR product bands were quantified using ImageJ and shown as included exon percentages (inclusion band intensity/total intensity) in the bar plot below. A chi-squared test followed by multiple comparison testing was used to compare WT with experimental groups. ***, p < 0.001. ns, p > 0.05. (E) Venn diagram of overlapping SE events between WT vs U2-2^WT/C28T^, WT vs U2-2^KO/KO^, and WT vs U2-2^C28T/KO^ comparisons. (F) Venn diagram of overlapping RI events between WT vs U2-2^WT/C28T^, WT vs U2-2^KO/KO^, and WT vs U2-2^C28T/KO^ comparisons.

Given the known correlation between splicing and gene expression, we next examined gene expression changes resulting from U2-2 snRNA dysregulation. Principal component analysis (PCA) based on the gene expression profiles shows the replicates cluster by genotype (Figure 3A). PCA2 distinguished clusters by U2-2 copy number; U2-2 WT exhibited the highest U2-2 expression—showing distinct gene expression patterns relative to heterozygous and KO conditions of ∼2,000 highly variable genes across different U2-2 genotypes (Figure 3B). Differential expression analysis identified 2,067 upregulated genes and 2,138 downregulated genes in U2-2 WT versus U2-2^KO/KO^ (Figure 3C); 1,002 upregulated genes and 1,264 downregulated genes in WT versus U2-2^WT/C28T^ (Figure 3D); 1002 upregulated genes and downregulated 1264 genes in U2-2 WT compared to the U2-2^C28T/KO^ (Figure 3E). Notably, these DEGs from different comparisons have many intersecting genes (Figure 3F). GO analysis reveals that functional pathways, particularly RNP complex biogenesis, ribosome biogenesis and rRNA processing are consistently impacted (Figure 3C-E, lower panels). Together, these findings demonstrate that U2-2 impacts a broad range of genes, many of which are involved in RNA processing pathways. This further suggests that the loss or disruption of U2-2 may alter splicing regulation through secondary targets.

**Figure 3.**
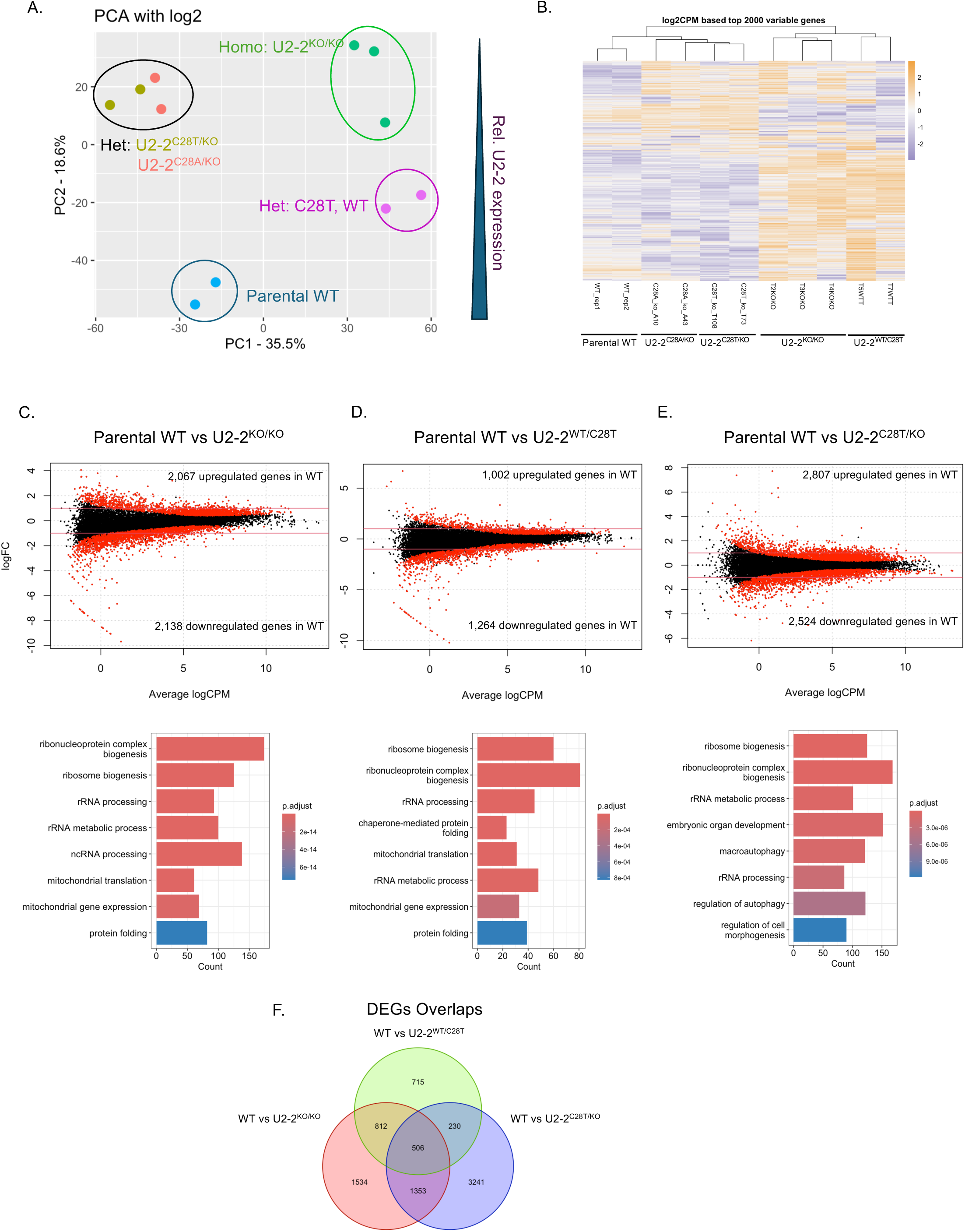
Differential expression analysis of U2-2 KO and mutants of HEK293T cells. (A) Principal component analysis (PCA) of transcriptomes from parental WT and U2-2 KO/mutant HEK293T cells. PC2 distinguishes samples by U2-2 expression status, and PC1 accounts for batch effects. (B) Heatmap showing the top 2000 variable genes across WT and mutant/KO samples, based on log_2_CPM values. (C-E) Top panels: Asymmetric changes in gene expression for comparing parental WT vs U2-2 mutant conditions. Adjusted p value ≤ 0.05 | Fold change > 1. Parental WT replicates n=3; U2-2 mutants replicates n=2. Low panels corresponding to the top panels: Gene Ontology (GO) analysis of differentially expressed genes reveals enrichment in functional pathways. (F) The Venn diagram of DEG overlapping genes between WT vs U2-2^WT/C28T^, WT vs U2-2^KO/KO^, and WT vs U2-2^C28T/KO^ comparisons.

### Absence of U2-2 snRNA predominantly leads to downregulation of genes with altered splicing

Given that U2-2 dysregulation impacts both splicing and gene expression, we analyzed the intersection of genes affected by these two processes to gain insights into potential downstream consequences of U2-2 activity. 477 genes (in red) exhibited both differential expression and altered splicing, suggesting that U2-2 snRNA influences both the transcriptional and post-transcriptional regulation of these genes (Figure 4A). An alternative visualization of this result indicates that 477 of the 4,275 differentially expressed genes also showed splicing alterations (Figure 4B). Further, pathway enrichment analysis reveals that these 477 genes are significantly associated with ribonucleoprotein complex biogenesis, ribosome biogenesis, and RNA splicing, underscoring the role of U2-2 in RNA processing and protein synthesis. More importantly, a subset of 377 genes from this group displayed downregulated expression in the U2-2 KO cell lines, with enrichment in the same pathways. This overlap highlights the extent to which U2-2 may be integral to the maintenance of cellular RNA and ribosomal machinery. Thus, U2-2 may serve as a regulatory node, with its disruption potentially compromising RNA splicing and ribosomal machinery.

**Figure 4.**
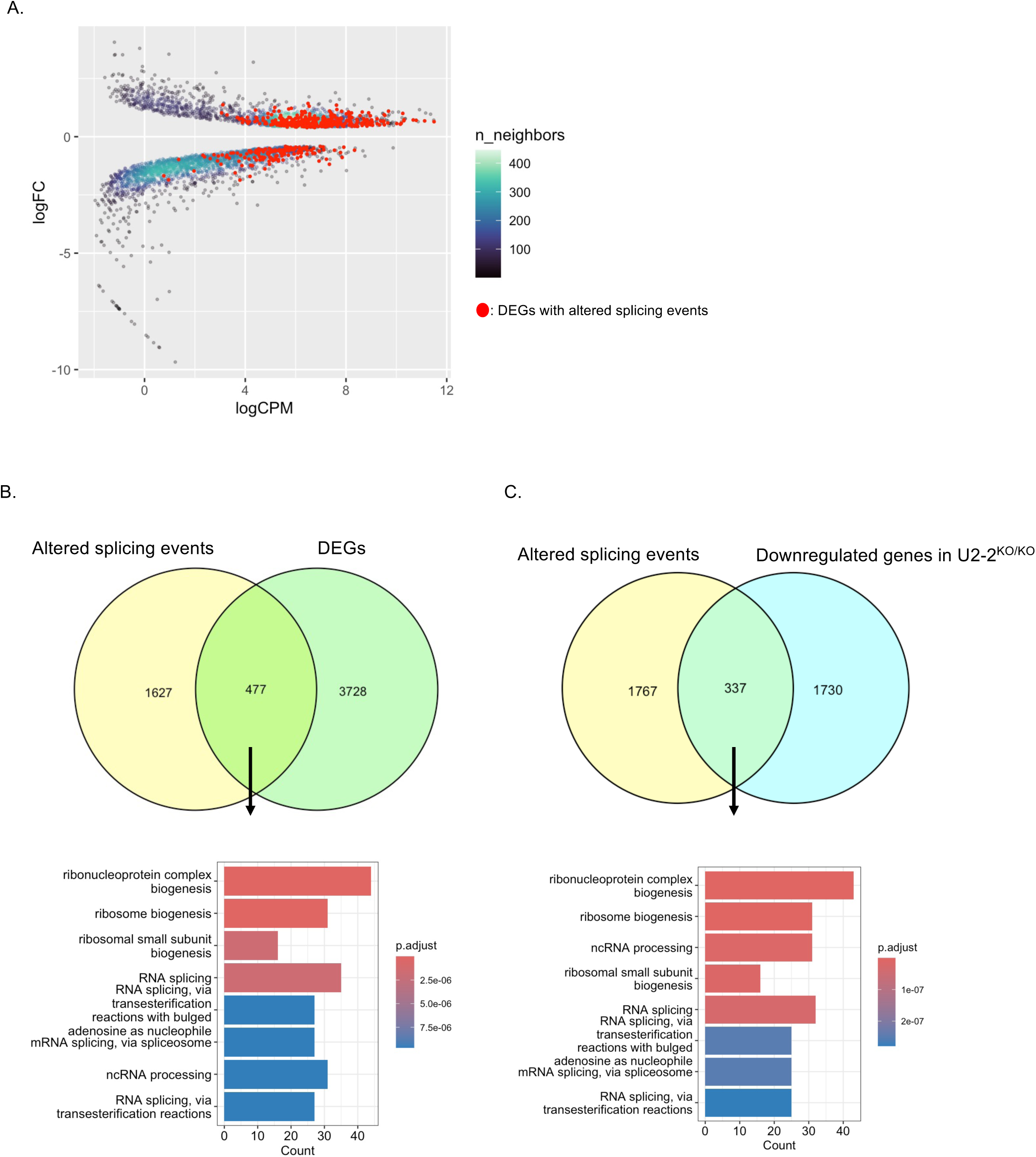
Analysis of alternative splicing events and differential genes upon knocking out U2-2 in HEK293T. (A) Scatter plot of overlaps between genes with altered splice events and DEGs comparing parental WT vs U2-2 KO, with altered splicing gene density on the right. Red dots represent differential expressed genes with altered splicing events. (B) Top: Venn diagram illustrating the intersecting of altered splicing events and DEGs comparing WT vs U2-2 KO. Lower: Enrichment pathways from intersecting of altered splicing events and DEGs comparing WT vs U2-2 KO. (C) Top: Venn diagram illustrating the intersecting of altered splicing events and downregulated genes in U2-2 KO groups comparing to the WT. Lower: Enrichment pathways from intersecting of altered splicing events and downregulated genes in U2-2 KO groups comparing to the WT.

### Overexpression of U2-2 WT and C28T mutant alters gene expression profiles and splicing

Because the C28T mutant expressed a compound phenotype (mutation + reduced U2-2 levels), we expressed the mutant in a background of WT level of WT U2-2 to avoid – to the extent possible – the effects of altered U2-2 levels. We expressed U2-2 WT and C28T mutant in parental HEK293T cells using lentiviral transfection (Figure 5A and Supplemental Figure S4A). Lentiviruses were produced using HEK293T cells carrying U2-2 WT, C28T mutant or empty vector (EV) constructs and subsequently infecting HEK293T destination cells. Approximately 1.5 × 10⁵ HEK293T cells produced a specific quantity of viruses (black box) that restores U2-2 expression in U2-2^KO/KO^ compared to parental HEK293T levels (Supplemental Figure S4B). Thus, U2-2^KO/KO^ destination cells were infected with the determined number of viruses carrying U2-2 WT, C28T, or empty vector constructs. Furthermore, the OE of U2-2 C28T mutant led to changes in gene expression and splicing, as revealed by RNA sequencing. Principal component analysis (PCA) (Figure 5B) and a heatmap of differential expression (Figure 5C) demonstrate clear separation between the U2-2 WT OE and C28T OE mutant compared to empty vector groups, indicating distinct transcriptional profiles based on U2-2 expression levels. Additionally, 3,234 genes were significantly upregulated, and 2,938 genes were significantly downregulated in U2-2 WT OE compared to the EV control. Additionally, 2,272 genes were significantly upregulated, and 1,943 genes were significantly downregulated in U2-2 C28T OE compared to empty vector control. In contrast, there were only 2 and 32 genes significantly upregulated and downregulated, respectively, in U2-2 WT compared to C28T OE. DEGs comparing WT OE vs empty vector with altered splicing events are mainly downregulated genes (207 out of 366 genes) (Venn diagrams, Figure 5D). These intersecting genes belong to pathways enriched for RNA splicing (Figure 5E).

**Figure 5.**
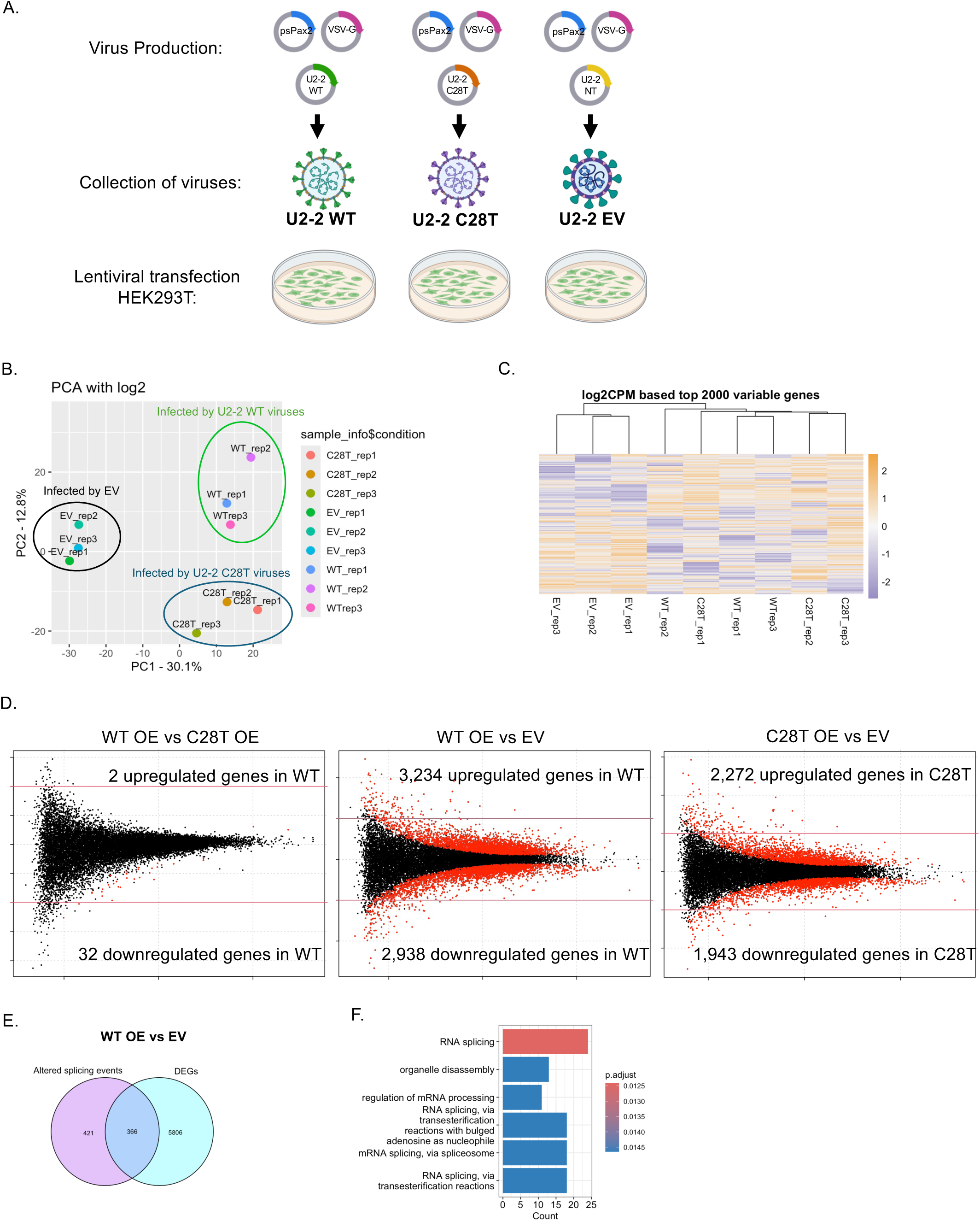
Overexpression of U2-2 WT and C28T mutant alters gene expression of HEK293T cells. (A) Workflow of lentiviral transfection to overexpress U2-2 WT, C28T mutant, and empty vector in HEK293T cells. (B) PCA of gene expression in cells overexpressing U2-2 WT, C28T mutant, and empty vector. PC1 showing clustering based on U2-2 expression levels. n=3. (C) Heatmap showing the top 2000 variable genes across of U2-2 WT OE, C28T OE mutant and empty vector replicates (n=3) based on log_2_CPM values. (D) Asymmetric changes in gene expression for comparing between three different overexpressing groups. p value ≤ 0.05 | Fold change > 1. (E) Venn diagram of the intersection of DEGs and altered splicing events between U2-2 WT OE vs empty vector. (F) Pathways of intersecting genes of DEGs and altered splicing events between U2-2 WT OE vs empty vector.

The greater impact of U2-2 snRNA dosage on DEGs compared to the U2-2 C28T mutation was further confirmed by overexpression. OE of U2-2 WT and the C28T mutant vs empty vector also showed a greater number of splicing alterations than U2-2 WT OE vs C28T OE conditions (Supplemental Figure S4A and S4B). Together, these findings suggest that U2-2 dosage, rather than mutation alone, plays a pivotal role in regulating splicing patterns, and the absence of U2-2 disrupts core components of the splicing pathway.

### Altered splicing due to the absence of U2-2 snRNA leads to nonsense-mediated decay (NMD)

NMD plays a role in the quality control of gene expression (Kurosaki et al., 2019), identifying and degrading transcripts that contain stop codons prior to the last exon junction complex (so-called premature termination codons, PTCs). Such PTCs may be caused by mutations or splicing defects, or more commonly are part of feedback circuits to regulate gene expression. Over 30% of SE events and 80% of RI events due to the absence of U2-2 snRNA have PTCs that lead to predicted NMD (Figure 6A). To determine the relationship between altered splicing due to U2-2 snRNA and NMD, we treated U2-2^KO/KO^ HEK293T cells with the ribosome-inhibiting drug cycloheximide (CHX) for 6 hours to block NMD, using dimethyl sulfoxide (DMSO) as the control. CHX treatment has been previously reported to increase detection of poison exon 6A in the *HNRNPL* gene (Rossbach et al., 2009), which we used as a positive control (Figure 6B). As expected, exon 6A-containing transcripts were detected in RNA from the CHX-treated WT and U2-2^KO/KO^ HEK293T cells but not in RNA from DMSO-treated cells, validating that CHX inhibited NMD in our experimental conditions. In RNA-seq analysis of these RNAs, 2,566 SE events were detected comparing the WT to U2-2^KO/KO^ (Figure 6C), similar to the extent of splicing alterations resulting from U2-2 depletion shown in Figure 2A. In contrast, a total of 6,106 SE events were identified when comparing DMSO-treated U2-2^KO/KO^ cells to CHX-treated U2-2^KO/KO^ cells (Figure 6C), indicating the significant portion of altered-splice events that are degraded by NMD in control cells. For example, the exon and intron inclusion rates of three example genes—*LUC7L*, *SNRNP70*, and *ZNF207*—are 3-4-fold higher in CHX-treated U2-2^KO/KO^ cells compared to DMSO-treated (Figure 6D). *LUC7L* is a known target of NMD that contains an autoregulatory poison exon (Choudhary & Norris, 2024; Mudge et al., 2011; Tan et al., 2020). *SNRNP70* exhibits both increased exon inclusion and increased intron retention (essentially an altered branch-3’ss use) that includes many stop codons, and the *ZNF207* exon is alternatively spliced in EMT (Pradella et al., 2017). We conclude that that many of the U2-2 KO altered mRNAs are degraded by NMD, and that the NMD-inhibited condition may provide a more accurate view of the U2-2 KO effects on the transcriptome.

**Figure 6.**
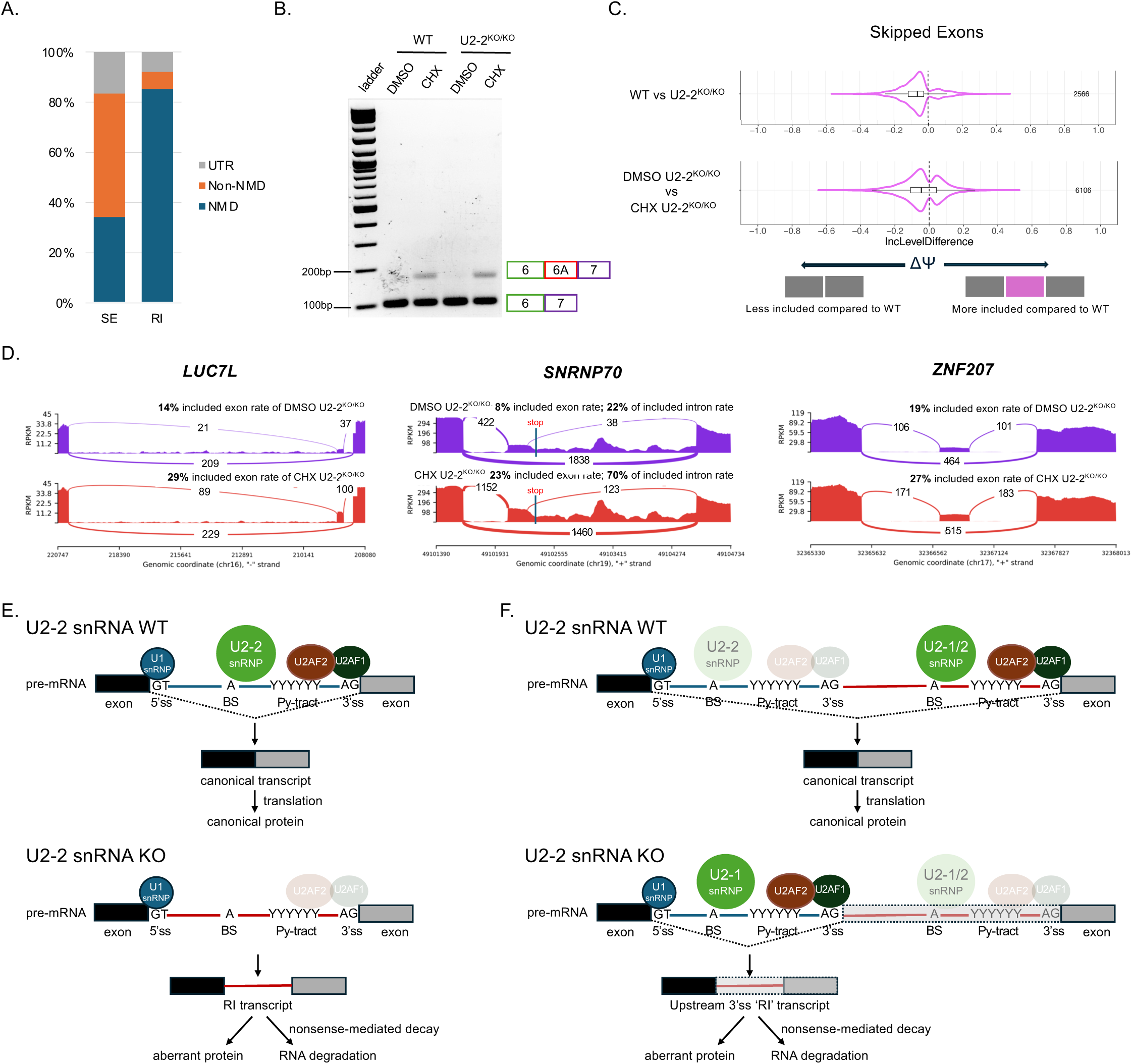
Inactivation of NMD enhances visualization of altered splicing. (A) A graph illustrating the percentage of alternatively spliced isoforms, including SE and RI, comparing WT vs U2-2 KO cells, categorized based on their outcomes: NMD, non-NMD and changes in UTR, predicted according to the PTC algorithm by Gupta, V. (Weinreb et al., 2022). (B) Exon 6A of the human *HNRNPL* gene serves as a positive control for CHX treatment (6 hours). Total RNA from WT and U2-2 KO cells was analyzed for exon 6A inclusion by RT-PCR, with schematic representations of expected RT-PCR products shown on the right. (C) Comparison of ΔΨ of SE events in WT vs U2-2^KO/KO^ (part of the Figure 2A) and DMSO U2-2^KO/KO^ vs CHX U2-2^KO/KO^. p ≤ 0.05 cut-off. The total number of significant altered events are shown on the right. (D) Sashimi plots of altered SE events of *LUC7L*, S*NRNP70* and *ZNF207* in different conditions through RNA-sequencing results. The RPKMs for each splice site are shown by the numbers overlying the arcs. The purple sashimi plots are SE in DMSO U2-2^KO/KO^ for each gene; the red sashimi plots are SE in CHX U2-2^KO/KO^ for each gene. (E) At some introns, U2-2 KO results in retained introns containing stop codons, which are sensitive to NMD. CHX treatment stalls ribosomes and results in significant enhancement of RI and SE signals, enhancing the sensitivity of the analysis. (E) At other introns, U2-2 KO results in increased use of upstream branch-3’ss, which, in the absence of U2-2 snRNPs, are recognized by U2-1 snRNPs. The included large intron regions contain multiple in-frame stop codons, leading to NMD. Increased exon inclusion (not shown) would be mechanistically similar.

## DISCUSSION

U2 snRNA is critical for formation and activity of the spliceosome and perhaps for other gene expression processes (van der Feltz & Hoskins, 2019). Several variant gene loci of U2 snRNA are expressed in humans, and mutations within highly conserved intron-recognition sequences are associated with cancers. So far, there have been no functional studies of these variants or mutants. We have generated U2-2 snRNA mutant and KO lines in HEK293T cells using CRISPR/Cas9, followed by RNA-sequencing and found that the U2-2 mutant has effects on gene expression and splicing patterns based both on the dosage of U2-2 and on the mutation per se.

Both the U2-2 mutant and KO lines of HEK293T exhibited altered gene expression as well as splicing patterns. However, our results suggest that U2-2 snRNA dosage exhibits a greater impact on gene expression and splicing than does the C28T heterozygous mutation of U2-2 (Figure 2 and 3). This finding indicates that the U2-2 mutation and dosage regulate the splicing through distinct regulatory mechanisms, and that the effects due to dosage can mask the effects due to mutation alone. The U2-2^KO/KO^ lines exhibited predominant downregulation of genes with altered splicing patterns: among 4,275 DEGs identified in U2-2 WT vs U2-2^KO/KO^ comparison, 477 genes also displayed splicing alterations (Figure 4B). Notably, these intersecting genes belong to enrichment pathways such as RNP complex biogenesis, ribosome biogenesis, and RNA splicing. In addition, U2-2 WT and C28T mutant were overexpressed in HEK293T cells to determine its effect using an alternative approach. U2-2 snRNA dosage had greater impact on DEGs and splicing alternations, as observed when comparing OE WT and OE C28T to empty vector in HEK293T cells (Figure 5 and Supplemental Figure S4). Lastly, our findings demonstrate that the majority of altered splicing events are predicted to trigger NMD (Figure 6A). Consistent with this, inhibition of the NMD pathway with CHX treatment resulted in an increased extent of altered splicing (Figure 6B-D).

### The U2-2 BSL mutant C28T affects splicing and gene expression

We used CRISPR-Cas9 cutting and homologous recombination to replace the U2-2 locus with mutant C28T in human embryonic cancer cell line HEK293T as heterozygous U2-2^WT/C28T^. C28 point mutations led to alterations in splicing (Figure 2F and 2G) and gene expression patterns (Figure 3A and 3B). There are 2,264 DEGs and many RNA splicing related pathways affected due to C28T point mutation, such as ribosome biogenesis, RNP complex biogenesis and rRNA processing etc. These results suggest that U2-2 mutation C28T within the BSL exerts its effects on splicing through perturbation of an interconnected network of genes. These genes, some of which could be influenced as secondary targets of the mutation (e.g. by altered levels of other splicing factors), contribute to widespread transcriptomic dysregulation.

### U2-2 dosage effects the transcriptome more than mutant alone

Both U2-2 snRNA mutant and KO lines were compared to WT HEK293T cells. Not only did point mutation C28T affect splicing and gene expression, U2-2^KO/KO^ and U2-2^C28T/KO^ lines exhibited larger effects than the point mutation itself, specifically the number of DEGs in WT vs U2-2^KO/KO^ and U2-2^C28T/KO^ (Figure 3C and 3E). These numbers are larger than comparing WT vs U2-2^WT/C28T^ (Figure 3D). Additionally, the U2-2 C28T mutant exhibits only 34 significant DEGs compared to WT (Figure 5D), using overexpression in HEK293T cells. In contrast, the RNA-seq data revealed a dramatic increase in DEG counts when comparing WT OE to empty vector controls (6,172 DEGs, Figure 5E) and C28T OE to empty vector controls (4,215 DEGs, Figure 5F).

Furthermore, 337 out of 477 of DEGs in the absence of U2-2 snRNA are downregulated genes with altered splicing (Figure 4B and 4C). This finding highlights the interplay between transcriptional regulation and splicing alterations due to the level of U2-2 snRNA. Different snRNAs have consistent roles in constitutive and alternative splicing, genes are particularly sensitive to variations in snRNA levels tend to be preferentially mis-spliced within a clinically diverse cohort of invasive breast ductal carcinomas (Dvinge et al., 2019). For instance, previous studies have reported that U11 snRNA alters gene splicing events and gene expression by affecting multiple signaling pathways in T24-related cell lines (Wang et al., 2021). U2 snRNP plays a crucial role in facilitating the efficient transition of promoter-proximal paused RNA polymerase II into productive transcription elongation (Caizzi et al., 2021). This finding highlights that U2-2 snRNA dosage implicates the regulations of splicing and transcription machinery. Therefore, our results demonstrate the sensitivity of transcriptomes to the level of U2-2 snRNA, emphasizing the complex and dosage-dependent regulatory functions of U2-2 snRNA.

DEGs from three comparisons (WT vs U2-2^KO/KO^, U2-2^C28T/KO^ and U2-2^WT/C28T^), including DEGs with altered splicing events, all belong to similar enrichment pathways (Figure 3, 4 and 5). In particular, RNP complex biogenesis, ribosome biogenesis, RNA processing and rRNA processing pathways were impacted by the dosage of U2-2 snRNA. Cajal bodies (CBs), also known as coiled bodies, are a subnuclear domain in the eukaryotic nucleus that contains proteins and factors involved in RNP biogenesis, rRNA precursor processing, U2 snRNA modification, and possibly post-transcriptional and detained-intron splicing (Hebert, 2010, 2013). The absence of U2-2 snRNA may disrupt the regulation of RNA processing factors associated with these identified gene pathways, impacting splicing in CBs.

### U2-2-dependent splicing changes and NMD

NMD is an RNA surveillance mechanism that identifies and degrades mRNA transcripts containing PTCs. This contributes to at least two distinct cellular process (Kurosaki & Maquat, 2016): quality control of gene expression and regulation of physiological mRNA levels. NMD identifies and degrades abnormal transcripts arising from routine errors in gene expression that introduce PTCs. NMD also contributes to gene expression homeostasis by selectively downregulating specific physiological mRNAs in response to cellular conditions. In our U2 KO studies, we showed altered splicing and gene expression patterns compared to the WT. Notably, the majority of SE and RI events resulting from splicing defects due to U2-2 absence generate PTCs, leading to predicted NMD activation (Figure 6A). Inhibition of the NMD pathway using CHX treatment further enhanced the detection of altered splicing (Figure 6C and 6D), providing a truer view of the extent of altered splicing, without bias by NMD. This reinforces the conclusion that U2-2 snRNA depletion disrupts selected splice events.

What is the basis for this selectivity? Are there binding sites for U2-2 snRNPs that are distinct from those of U2-1 snRNPs? If so, how is that mediated, as the branch recognition sequences (GUAGUA), and even the entire BSLs, are identical in U2-1 and U2-2 snRNAs? For some introns, U2-2 KO resulted in loss of splicing (Figure 6E). In other cases, the increased use of many branch-3’ss (resulting in increased exon inclusion or increased alternative branch-3’ss, sometimes hundreds of nucleotides upstream of the constitutive branch-3’ss) in the U2-2 KO suggests that these sites are derepressed in the KO. Much future work will be needed to understand the mechanism of such derepression. Possibilities include that the binding of U2-2 snRNP or factors in its downstream network at these sites are inhibitory at the assembly stage, and one interpretation is that these are sites where U2-2 snRNPs have a higher affinity than U2-1 snRNPs, but in the absence of U2-2 snRNPs, U2-1 snRNPs bind and are more active in promoting splicing than are U2-2 snRNPs (Figure 6F). Alternatively, U2-2 spliceosomes in combination with certain splice site sequences may be more prone to discard/disassembly (e.g. recognized as ‘aberrant’ pre-spliceosomes by SUGP1 and disassembled by the RNA helicase Prp43/DHX15 or recognized as aberrant B^act^ spliceosomes and disassembled by DHX35– GPATCH1 and Prp43/DHX15–NTR1) (Beusch & Madhani, 2024; Li et al., 2025; Soni et al., 2025). Consistent with the former possibility, it is notable that there are far more U2 snRNPs than U4/5/6 snRNPs (∼500,000 and 200,000 per nucleus, respectively), and thus, like the even more abundant U1 snRNP (∼1,000,000 per nucleus), U2-2 snRNPs may have roles other than promoting productive splicing.

### Limitations of the study

Future studies should investigate the impact of U2-2 depletion and mutation across diverse cellular and tissue models to assess our findings beyond HEK293T cells. Furthermore, biochemical and structural analyses will be necessary to elucidate the molecular interactions between U2-2 mutant snRNA and other splicing factors. This would provide additional mechanistic insight into its role in splicing regulation. In addition, the effects of mutation within U2-1 should be investigated, as well as the role of U2-1 copy number on the penetrance of the mutant’s effects. Such questions are currently limited by the difficulty in manipulation of a multicopy gene locus.

## RESOURCE AVAILABILITY

### Lead contact

Further information and requests for resources and reagents should be directed to and will be fulfilled by the lead contact, Charles Query.

### Material availability

The plasmids and cell lines generated in this study are available upon request.

### Data and code availability

- All data reported in this paper will be shared by the lead contact upon request. All data supporting the findings of this study are available within the article and its supplemental information.
- This paper does not report original code.
- Any additional information required to re-analyze the data reported in this paper is available from the lead contact upon request.

## ACKNOWLEDGMENTS

We are grateful to Dr. Matthew Gamble, Dr. Teresa Bowman, Dr. Julie Secombe and Dr. Wenjun Guo for helpful discussions and to members of the Flow Cytometry Core Facility at Albert Einstein College of Medicine for expert assistance. This work was supported by NIH grant GM57829 to C.C.Q and Cancer Center Support Grant P30CA013330 from the NCI to AECOM/MECCC.

## AUTHOR CONTRIBUTIONS

Conceptualization, D.C., V.G. and C.Q.; Methodology and Investigation, D.C. and V.G.; Software, V.G.; Data Curation and Analysis, D.C. and V.G.; Writing – Original Draft, D.C. and C.Q.; Writing – Review & Editing, D.C., V.G. and C.Q.; Funding Acquisition and Supervision, C.Q.

## STAR METHODS

Detailed methods are provided in the online version of this paper and include the following:

- KEY RESOURCES TABLE
- EXPERIMENTAL MODEL AND STUDY PARTICIPANT DETAILS

`o` HEK293T embryonic kidney cells
- METHOD DETAILS

`o` Generation of HEK293T U2-2 KO and KI by single-cell cloning
`o` Generation of U2-2 OE lines
`o` Plasmids
`o` DNA and RNA isolation from cell lines
`o` RT-PCR
`o` Bioinformatic analysis of RNA-sequencing results
`o` Amplicon sequencing
- QUANTIFICATION AND STATISTICAL ANALYSIS

## SUPPLEMENTAL INFORMATION

Supplemental information can be found online at xxxx.

## EXPERIMENTAL MODEL AND SUBJECT DETAILS

### Cell cultures

HEK293T cells were a gift of Dr. Matthew Gamble (Albert Einstein College of Medicine) and validated by Cell Line Authentication (CLA) service at the Einstein Genomics Core. All cells were cultured in 90% DMEM, 10% FBS, and 1x Pen/Strep and grown at 37°C in 5.0% CO2 in a Heracell Vios 160i incubator.

## METHOD DETAILS

### Generation of HEK293T U2-2 KO and mutants by single-cell cloning

HEK293T cells were transiently transfected with two pSpCas9(BB)-2A-GFP (PX458) vectors containing sgRNAs targeting sequences upstream and downstream of the mature U2-2 snRNA (Kosmyna et al., 2020) using the jetOPTIMUS transfection reagent commercial kit (Polyplus). The knock-in lines were generated with additional template plasmids (GenScript) carrying the U2-2 mutation for the homologous recombination. After 48-72 hours, GFP-positive transfected cells were single-cell sorted into 96-well plates using FACS Moflo XDP sorting machine. DNA from clonal cells was isolated using Zymo Quick DNA miniprep with Zymo-spin IIC columns. PCR primers flanking the sgRNA targets were using for genotyping utilizing NEB Taq 2x Master Mix (M0207). sgRNA and primer sequences are listed in Supplemental Table S1. The genotyping sequences of 3 U2-2^KO/KO^ replicates are provided in Supplemental Table S2.

### Generation of U2-2 Overexpression (OE) lines

OE of U2-2 WT, C28T and empty vector were generated using the pLVX-puro lentiviral system (Clonetech) in HEK293T cells. pLVX-puro vectors were described in the Plasmids section, Lenti-PAX2 and VSV-G plasmids for viral production were transfected into HEK293T cells using jetOPTIMUS transfection reagent protocol (Polyplus). 3.2 ug Lenti-PKG, 1.2 ug VSV-G and 4 ug of pLVX-puro rescue vectors were mixed. This master mix was vortexed and spined down, then 2 ul of jetOPTIMUS reagent was added to the master mix. After 10 min incubation at RT, the master mix was added to HEK293T cells at approximately 40-50% confluency of a 35 mm plate. Viruses were collected for three days consecutively using 0.45 um filters and stored in a 4-degree fridge. Lenti-X^TM^ concentrator (Clonetech PT4421-2) was used to concentrate lentiviral stock. Destination cells were infected at approximately 60-70% confluency. After 24 hrs, the media was changed and the infected HEK293T cells were selected in media supplemented with 0.5 μg/mL puromycin for at least 3 passages. The negative controls (Parental HEK293T cells) were dead within 3 days in the selection media.

### Plasmids

Plasmids expressing guide RNAs used for CRISPR/Cas9 KO of the U2-2 locus were gifts from Dr. Brian Kosmyna (Kosmyna et al., 2020). Plasmids carrying U2-2 mutant templates used for KI were purchased from GenScript.

Plasmids used for U2-2 OE were pLVX-puro-dCMV and derivatives described below [from Dr. Brian Kosmyna (Kosmyna et al., 2020) and Lenti-PAX2 (Addgene #12260) and VSV-G (Addgene #138479). pLVX-puro-U2-2-WT and pLVX-puro-C28T were constructed from pLVX-puro lentiviral expression plasmid (Clonetech). pLVX-puro was digested with EcoRI (NEB 10227213). The insert containing U2-2 WT and C28T mutant were ligated into linearized pLVX-puro using NEB HiFi assembly kit. These constructs were confirmed by Sanger sequencing. PCR primer sequences for amplification of U2-2 are listed in Supplemental Table S1.

### DNA and RNA isolation from cell lines

Genomic DNA from cell lines was isolated using Zymo Quick DNA miniprep with Zymo-spin IIC columns (Catalog no. D3024). Total RNA was isolated from 3 consecutive passages of the control group parental cells and from clonal mutant cell lines of HEK293T using Trizol reagent (Invitrogen; catalog numbers 15596026 and 15596018) according to manufacturer’s protocol at approximately 90% confluency. RNA was quantified first on a Nanodrop2000 and diluted to 400 ng/ul. Final concentrations of RNA were then determined on a Denovix QFX fluorometer using a Qubit RNA BR Assay kit according to the manufacturer’s protocol.

### RT-PCR

RNA was isolated as described above. cDNA from 400 ng of total RNA was reverse transcribed using the SuperScript™ III One-Step RT-PCR System with Platinum™ *Taq* DNA Polymerase (Invitrogen 12574026) with 50 pmol of random hexamer primers and in a 20 ul reaction, according to manufacturer’s protocol (First-Strand cDNA Synthesis). Specific gene primers and 100X diluted cDNA were added to NEB Taq 2x Master Mix for 40 ul of PCR reaction. Primer sequences were designed to detect the spliced state of each transcript at the event of interest and are listed in Supplemental Table S1. Subsequently, PCR products of each sample were loaded onto a 2% agarose/TBE gel with ethidium bromide. The quantity of DNA was determined by the intensity of PCR bands using ImageJ (Schneider et al., 2012). The unpaired, two-tailed Student’s t-test was applied as described above.

### Bioinformatic analysis of RNA-sequencing results

RNA was submitted to Genewiz (Azenta) for sequencing. RNA was poly(A) selected and sequenced to a depth of 40 million reads per sample. Raw FASTQ files were obtained from Genewiz and was checked for quality of reads with FASTQC (version 0.11.4). The raw reads for both bulk-control and experiments were aligned with a splice aware aligner such as STAR (Version 2.4.2).

#### Alternative splicing analysis

The alternative splicing analysis of RNA-seq data was performed using rMATS software (v4.1.0) (Shen et al., 2014). The rMATS statistical model for paired replicates was used to determine the expression of alternative splicing events. The Likelihood Ratio Test was used to determine the p-value of differences in inclusion level between two groups. The resulting p-values were adjusted using the Benjamini and Hochberg procedure to control for false discovery rate (FDR). The analysis was done for 5 types of alternative splicing events: SE, RI, MXE, A3SS and A5SS. Differential splicing events with FDR p-value <0.05 were considered significant.

#### Differential gene expression and GO enrichment analysis

Differential gene expression analysis between two groups was performed using the edgeR R package (Robinson et al., 2009). Genes with low counts (CPM < 0.5) were excluded from analysis. The resulting p-values were adjusted using the Benjamini and Hochberg procedure to control for false discovery rate (FDR). The DEGs were then selected using FDR p-value < 0.05. GO pathway enrichment analysis of the DEGs was performed using the clusterProfiler R package (Yu et al., 2012), and significant enrichment was determined using p-value < 0.05.

### Amplicon sequencing

#### Deep amplicon sequencing of U2-2 DNA

U2-2 DNA was amplified using primers (hU2-2-115 and hU2-2-117, or hU2-2-116 and hU2-2-118) containing barcodes, with NEB Taq 2x Master Mix. The PCR products were resolved on a 2% agarose/TBE gel, and bands of expected sizes were isolated. DNA was then purified using the Zymo DNA Clean and Concentrator kit (Catalog Number: D4003T). Purified products were submitted to Genewiz for Amplicon-EZ sequencing (150-500 bp range).

#### Deep amplicon sequencing of U2-1 and U2-2 RNAs

Total RNA was isolated using Trizol (described above) and subsequently reverse transcribed (RT) using antisense oligo hU2-109 primer, which base-pairs equally to U2-1 and U2-2 and contains a unique 10-nt molecular identifier (UMI) and partial Illumina adapter. Each reaction (1 μL of 10 μM primer, 400 ng RNA, 1.2 μL of 5X AB buffer, and water to 6 μL total volume) was heated to 90°C for 3 minutes and gradually cooled to 48°C. SuperScript™ III One-Step RT-PCR System with Platinum™ Taq DNA Polymerase (Invitrogen 12574026) was used, adding 0.5 μL enzyme (diluted 1:50 in buffer), 5X dNTP Mix (dGTP, dATP, dTTP, dCTP), 3 μL of 1X AB buffer, and 0.5 μL of 5X RT buffer to each reaction, with incubation at 37°C for 5 minutes and 42°C for 30 minutes. The final reaction mixture was diluted to 100 μL, and 1 μL was used for PCR amplification using primers hU2-110 (primes equally on U2-1 and U2-2 cDNA for 2^nd^ strand synthesis) and pIlAd-Rev. The remaining steps of isolation, purification, and quantification were identical to those described for “Deep Amplicon Sequencing of U2-2 DNA.”

#### Amplicon sequencing analysis

Raw Amplicon-EZ fastq files were directly downloaded from the GENEWIZ sFTP server. They typically provide ∼50,000 reads per sample. Bioconda cutadapt package was used to select the reads with the correct adapter sequences:

For the forward sequencing read: 5’-ACACTCTTTCCCTACACGACGCTCTTCCGATCT -3’ For the reverse sequencing read: 5’-GACTGGAGTTCAGACGTGTGCTCTTCCGATCT -3’ The designed PCR primers were added to the 5’ end of the primers. Reads were mapped to a customized reference genome (in silico reference genome with U2-1 and U2-2 snRNAs) to identify variants.

**Supplemental Figure S1. Comparison of U2-1 and U2-2 snRNAs and modulation of RNA structures in the spliceosome**

(A) Clustal Omega alignment (Sievers et al., 2011) of transcribed U2-1 and U2-2 snRNAs. Position C28 is circled in red.

(B) Secondary structure of the evolutionarily conserved BSL of U2 snRNA. Position C28 is circled in red. Base pairing interactions for Watson Crick interactions (–) or wobble interactions (•) are shown.

(C) Position C28 in the context of RNA-RNA interactions within the activated spliceosome. The pre-mRNA is shown in black, U2 snRNA is shown in dark red, U5 in grey and U6 in green as C complex. BS-U2 duplex site (with the branch site (BS) adenosine highlighted in blue). Base pairing interactions for Watson Crick interactions (–) or wobble interactions (•) are shown.

**Supplemental Figure S2. DpnII restriction fragment length polymorphism (RFLP) and genotyping of U2-2 mutant lines**

(A) Schematic of U2-2 RFLP. DpnII digested WT U2-2 gives three fragments, DpnII digested mutant U2-2 gives two fragments, and U2-2 KO is not digested by DpnII.

(B) Agarose gels of U2-2^WT/C28T^, U2-2^C28A/KO^, and U2-2^C28T/KO^ replicates. DpnII undigested and digested bands with sanger sequencing results.

**Supplemental Figure S3. Splice events are altered comparing WT to U2-2 KO and mutants of HEK293T**

(A) Schematics of five alternative splicing events in U2-2 WT vs different mutant conditions. Grey boxes indicate exons and lines indicate introns. Colorful boxes indicate mis-spliced exons corresponding to the splicing events.

(B) Comparison of ΔΨ of MXE events between WT vs three U2-2^KO/KO^, two U2-2^WT/C28T^, and two U2-2^C28T/KO^ cell lines at p ≤ 0.05 cut-off. The total number of significant altered events are shown on the right. Grey boxes indicate exons and lines indicate introns. Purple box indicates mis-spliced exons corresponding to the SE event.

(C) Comparison of ΔΨ of A5SS events between WT vs three U2-2^KO/KO^, two U2-2^WT/C28T^, and two U2-2^C28T/KO^ cell lines at p ≤ 0.05 cut-off.

(D) Comparison of ΔΨ of A3SS events between WT vs three U2-2^KO/KO^, two U2-2^WT/C28T^, and two U2-2^C28T/KO^ cell lines at p ≤ 0.05 cut-off.

**Supplemental Figure S4. RNA Amplicon deep sequencing results of U2-2^KO/KO^ destination cells by U2-2 viruses and the splicing alteration in SE and RI of the OE system**

(A) The y-axis represents the fractional ratio of U2-1 (blue) to U2-2 (orange) across various samples displayed on the x-axis. Ratios for HEK293T parental WT and U2-2^WT/C28T^ endogenous KI samples are shown for comparison with OE samples. Samples infected with U2-2 WT viruses are highlighted in black boxes, those infected with U2-2 C28T viruses are in green boxes and samples infected with empty vector are marked with red boxes.

(B) The number of U2-2 viruses required to recapitulate the U2-2 level of parental HEK293T cells was determined through a titration. The y-axis represents the fractional ratio of U2-1 (blue) to U2-2 (orange) across samples. The x-axis represents HEK293T parental WT compared to samples infected with U2-2 WT viruses produced at varying titration levels of HEK293T cells.

(C) Comparisons of ΔΨ of SE between U2-2 WT OE vs EV, WT OE vs C28T OE, and C28T OE vs empty vector at p ≤ 0.05 cut-off. The total number of significant altered events are shown on the right.

(D) Comparisons of ΔΨ of RI.

