## Supplemental Materials for "U2-2 snRNA Mutations Alter the Transcriptome"

### Supplemental Figure S1

A.

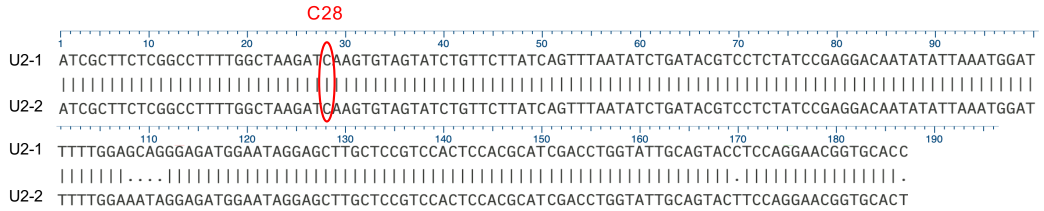


B.

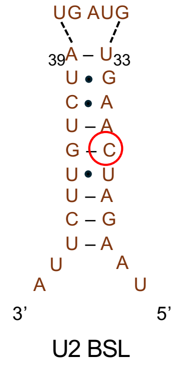


C.

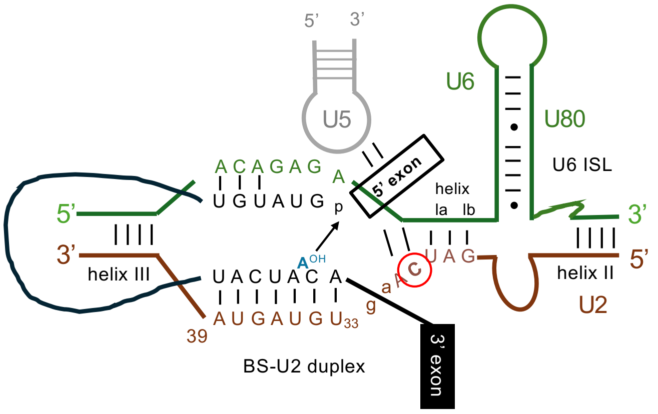

### Supplemental Figure S2

A.

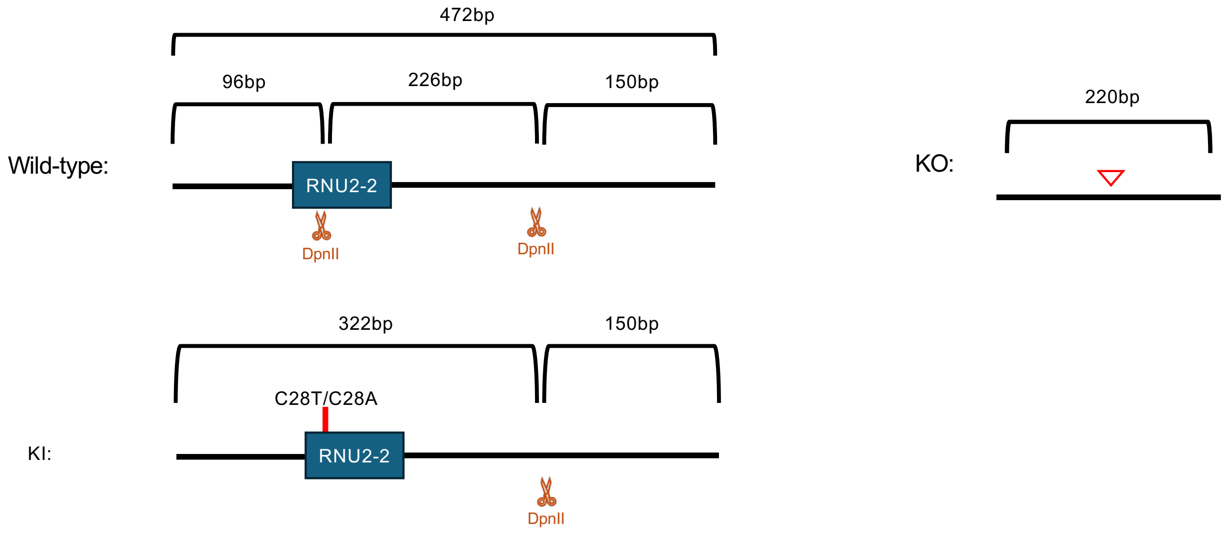


B.

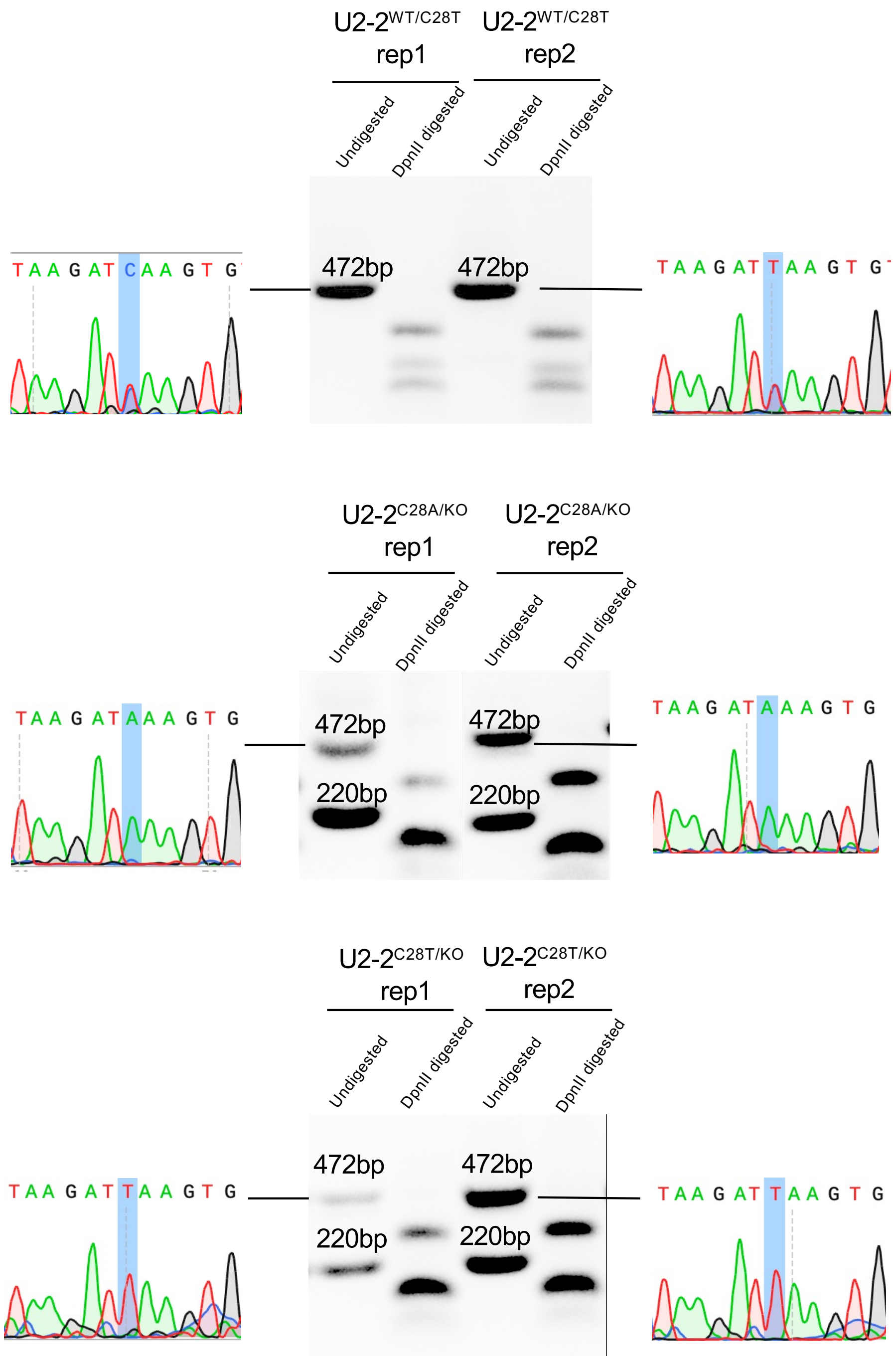

### Supplemental Figure S3

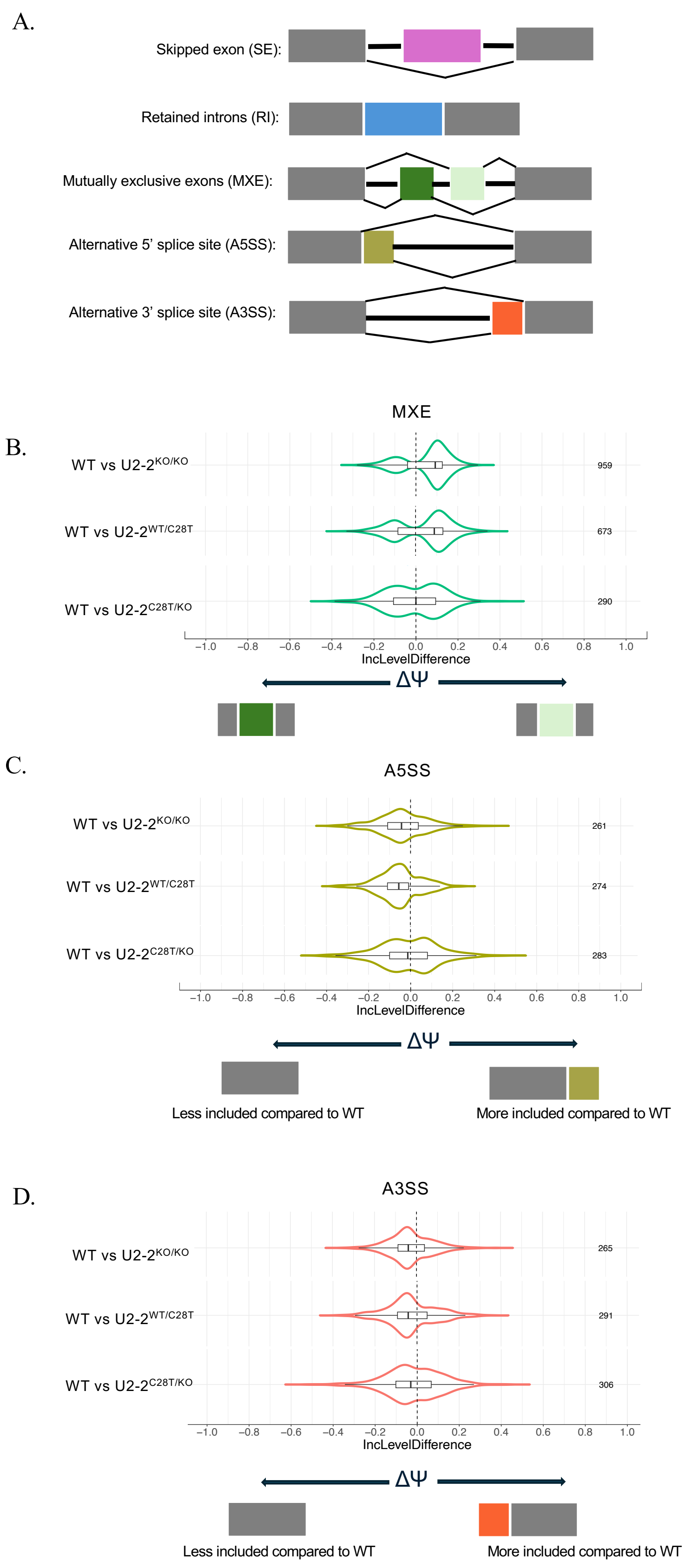

### Supplemental Figure S4

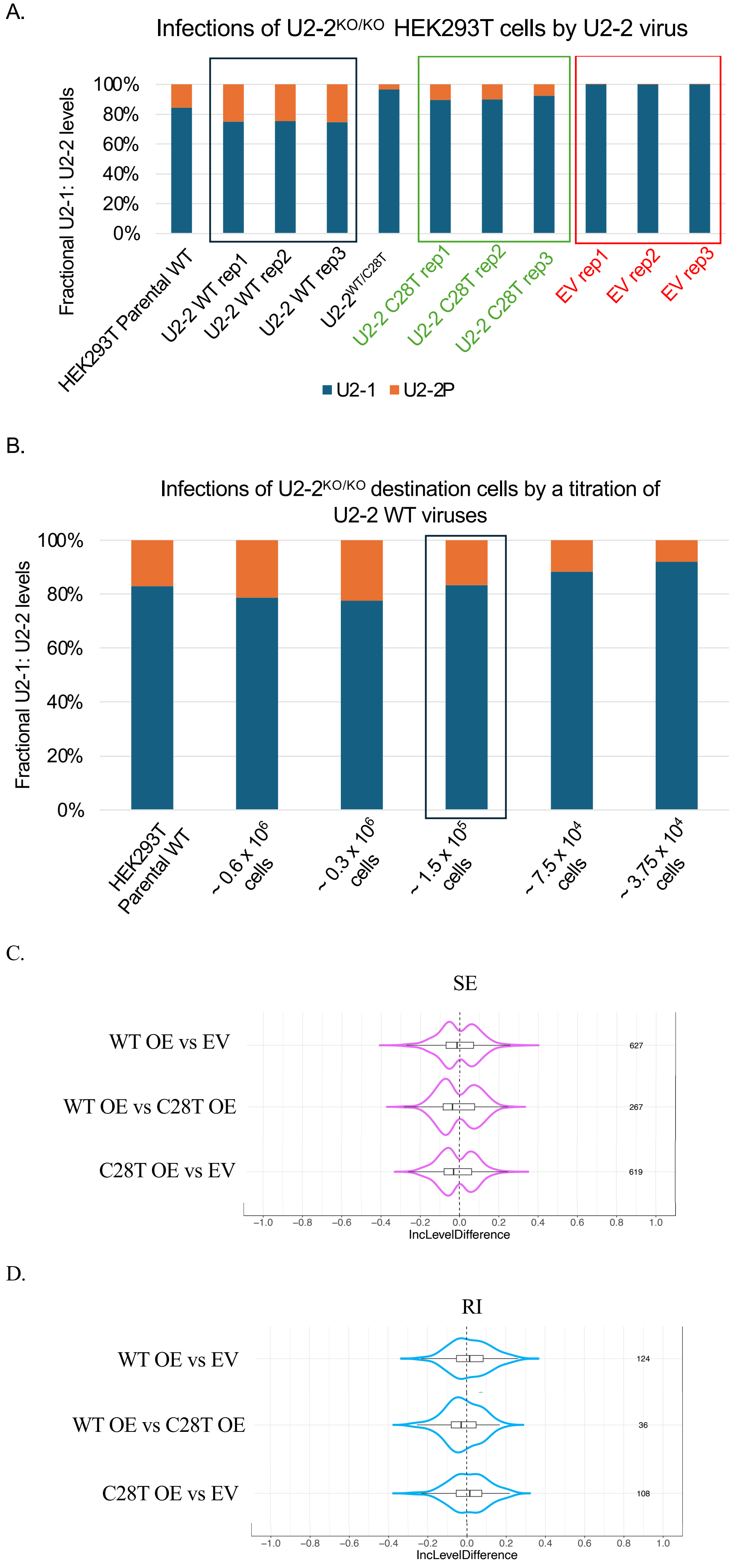

#### Supplemental Table S1: Primers used for genotyping and RT-PCR

| **Name** | **Sequence** (5' - 3') | **Notes** |
|---|---|---|
| gU2-2F | CCT GGA TAG TTA CCA TAA CTG G | Genomic U2-2 Forward |
| gU2-2R | GGA GAT ACT ACG CTC CGT GCT CAG | Genomic U2-2 Reverse |
| Luc7L Forward | CCCTCCGAGCAGATTATGAG | Genomic Luc7 Forward |
| Luc7L Reverse | GCAGAAACTTCCGCACTGAT | Genomic Luc7 Reverse |
| SnRNP70 Forward | CAGTAAGCGGTCAGGAAAGC | Genomic SnRNP70 Forward |
| SnRNP70 Reverse | TCCAGCCCTTCACGGTTC | Genomic SnRNP70 Reverse |
| ZNF207 Forward | ATAGACCACCTGCACCAACA | Genomic ZNF207 Forward |
| ZNF207 Reverse | GCAGGGAATGTAGGCTTTGG | Genomic ZNF207 Reverse |
| pIIAd-Rev | GACTGGAGTTCAGACGTGTGCTCTTCCGATCT | Partial Illumina adapter-reverse |
| hU2-1-106 | ACACTCTTTCCCTACACGACGCTCTTCCGATCT GGA CGA GAC AGA GGG AAT GG | IIAd+101 (for RNU2-1 amp-seq) |
| hU2-1-107 | GACTGGAGTTCAGACGTGTGCTCTTCCGATCT CCG CAC ATC AGG AAC CTC AAG | IIAd+105 (for RNU2-1 amp-seq) |
| hU2-109 | GACTGGAGTTCAGACGTGTGCTCTTCCGATCT NNNNNNNNNNTT TGG AGT GGA CGG AGC AAG CTC C | IIAd+BK125, contains UMI (for RT; primes equally on U2-1 and U2-2 for 1st strand synthesis) |
| hU2-110 | ACACTCTTTCCCTACACGACGCTCTTCCGATCT TTCC ATC GCT TCT CGG CCT TTT GG | IIAd+A+U2-5'end (primes equally on U2-1 and U2-2 cDNA for 2nd strand synthesis) |
| hU2-2-115 | ACACTCTTTCCCTACACGACGCTCTTCCGATCT GACT CCT GGA TAG TTA CCA TAA CTG G | IIAd+A+gU2-2F(A) |
| hU2-2-116 | ACACTCTTTCCCTACACGACGCTCTTCCGATCT TTGA CCT GGA TAG TTA CCA TAA CTG G | IIAd+B+gU2-2F(B) |
| hU2-2-117 | GACTGGAGTTCAGACGTGTGCTCTTCCGATCT GACT GGA GAT ACT ACG CTC CGT GCT CAG | IIAd+gU2-2R(A) |
| hU2-2-118 | GACTGGAGTTCAGACGTGTGCTCTTCCGATCT TTGA GGA GAT ACT ACG CTC CGT GCT CAG | IIAd+gU2-2R(B) |
| U2-2 CRISPR 5'-F | CACCGCGCCGAGGCGACCGAAGTAA | sgRNA |
| U2-2 CRISPR 3'-F | CACCGCGAAATCTTCCATTAAACAA | sgRNA |
| hnRNP L Forward | TAC ACA AAC CCC AAT CTC AGT | Genomic hnRNP L Forward |
| hnRNP L Reverse | TAT TCT GCG GGG TGA TCT CC | Genomic hnRNP L Reverse |

#### Supplemental Table S2: Genotyping results of 3 biological replicates of U2-$2^{KO/KO}$ of HEK293T cells

<table>
<tr><td>U2-$2^{KO/KO}$ rep1 sequence:<br><br>GCGGTATTGCAGTACGCCGCCGAGGCGACCGAA|GTTTAATGGAAGATTTCGATCAGTTA<br>GGGTACAAGCTAAATAGTTATGTTCTTGTTGTTTGAGTTGGATTAGGTGTTTTTATCGCTA<br>GTGTGTTGAATGACGCGTGTTTGAGTGTCTTGTTGAGAGAAGGTTTTGTTTTATAACTGA<br>GCACGGAGCGTAGTATCTCCC</td></tr>
<tr><td>U2-$2^{KO/KO}$ rep2 sequence:<br><br>CCAAGGTTGGCAGGTAGGCCGCCGAGGCGACCGA|GTTTAATGGAAGATTTCGATCAGT<br>TAGGGTACAAGCTAAATAGTTATGTTCTTGTTGTTTGAGTTGGATTAGGTGTTTTTATCGC<br>TAGTGTGTTGAATGACACGTGTTTGAGTGTCTTGTTGAGAGAAGGTTTTGTTTTATAACTG<br>AGCACGGAGCGTAGTATCTCCCCA</td></tr>
<tr><td>U2-$2^{KO/KO}$ rep3 sequence:<br><br>ACATTAACCAGAAGGCGCCGAGGCGACCGA|GTTTAATGGAAGATTTCGATCAGTTAGG<br>GTACAAGCTAAATAGTTATGTTCTTGTTGTTTGAGTTGGATTAGGTGTTTTTATCGCTAGT<br>GTGTTGAATGACGCGTGTTTGAGTGTCTTGTTGAAAGAAGGTTTTGTTTTATAACTGAGC<br>ACGGAGCGTAGTATCTCCAA</td></tr>
</table>

| : cut/repair site
